# Potential Key Genes Associated with Stroke types and its subtypes: A Computational Approach

**DOI:** 10.1101/2021.09.13.460051

**Authors:** Gourab Das, Pradeep Kumar

## Abstract

To investigate prospective key genes and pathways associated with the pathogenesis and prognosis of stroke types along with subtypes. Human genes using genome assembly build 38 patch release 13 with known gene symbols through NCBI gene database (https://www.ncbi.nlm.nih.gov/gene) were fetched. PubMed advanced queries were constructed using stroke-related keywords and associations were calculated using Normalized pointwise mutual information (nPMI) between each gene symbol and queries. Genes related with stroke risk within their types and subtypes were investigated in order to discover genetic markers to predict individuals who are at the risk of developing stroke with their subtypes. A total of 2,785 (9.4%) genes were found to be linked to the risk of stroke. Based on stroke types, 1,287 (46.2%) and 376 (13.5%) genes were found to be related with IS and HS respectively. Further stratification of IS based on TOAST classification, 86 (6.6%) genes were confined to Large artery atherosclerosis; 131 (10.1%) and 130 (10%) genes were related with the risk of small vessel disease and Cardioembolism subtypes of IS. Besides, a prognostic panel of 9 genes signature consisting of CYP4A11, ALOX5P, NOTCH, NINJ2, FGB, MTHFR, PDE4D, HDAC9, and ZHFX3 can be treated as a diagnostic marker to predict individuals who are at the risk of developing stroke with their subtypes.

## Introduction

Stroke is a complex heterogeneous disorder that occurs due to the interaction between environmental and genetic risk factors.[1, 2] It is one of the main important causes of mortality and long-term disability worldwide.[3] About 85% of stroke cases are ischemic stroke (IS), whereas 15% are hemorrhagic stroke (HS).[4] According to the Trial of Org 10172 in Acute Stroke Treatment (TOAST) classification; IS has been categorized according to the presumed etiological mechanism into five groups: large artery atherosclerosis (LAA), small vessel disease (SVD), cardio-embolic disease (CE), other determined etiology (ODE), and undetermined etiology (UDE).[5] Furthermore, HS is categorized into intracerebral hemorrhage (ICH) and Subarachnoid hemorrhage (SAH). Despite recent advancements in treatment modality, very few are known regarding the essential pathophysiology of stroke, and further research is still warranted to elucidate mechanisms in order to identify stroke occurrences.

Several established risk factors including diabetes, hypertension, dyslipidemia, smoking, atrial fibrillation, and obesity have been a link to the happening of stroke.[6] The fraction of strokes of undermined or rare causes is greater for young adults as compared to elders, and in many cases, underlying causes are genetic related. More than hundreds of genes have been described to be linked with the risk of stroke.[7, 8] Unravelling the genetic causes that play an important role in IS and HS is very challenging, as the genetic part of it is multifaceted.[9] In most cases, numerous genes are likely involved in the pathogenesis of stroke performing on a broad variety of candidate pathways, such as inflammatory, haemostatic, renin-angiotensin-aldosterone, and homocysteine metabolisms.[10, 11]

The genetic constituent is more predominant in LAA subtypes of IS than in SVD or cryptogenic IS and in patients younger than 70 years of age.[12] Previously published multicentric genetic studies using genome-wide data estimated that 40% for LAA, 33% for CE, 16% for SVD, and 38% for combined (Determined plus undetermined) etiology comprises the heritability of IS.[7, 13, 14] with the illustration that some genetic variants may serve as causal markers for stroke. To recognize the role of particular risk elements in regulating the pathophysiology of stroke, the hereditary basis of every risk factor is desirable to be examined and integrated, in context to their biological role and pathway interactions. To date, there are no well-established genetic markers that may discriminate the stroke types as well as their subtypes.

Identifying novel diagnostic and prognostic genetic markers has become an urgent demand. But its experimental determination remains a costly and time-consuming process. Hence, novel computational methods are needed to fulfil this requirement. But, very few *in silico* methods were developed in this regard including gene expression-based models [18], machine learning-based classifiers,[19] genetic algorithm-based models [20], and a relational database named SigCS base (http://sysbio.kribb.re.kr/sigcs) [21] which documented genes, variants, and pathways related to cerebral stroke. Unfortunately, this rich resource was discontinued as of February 2021. So, there was a huge scope for the development of a computational algorithm for the prediction of genes associated with stroke types and their sub-types. Our computational approach was aimed to recognize possible important genes and the pathways linked with the pathogenesis and prediction of stroke types along with their subtypes.

## Methods

### Advanced query building and searching PubMed

We fetched Human genes using genome assembly build 38 patch release 13 with known (status “Active”) gene symbols through the NCBI gene database (https://www.ncbi.nlm.nih.gov/gene). PubMed advanced queries were constructed using stroke-related keywords and associations were calculated using Normalized pointwise mutual information (nPMI) between each gene symbol and queries.[22] To reduce the false hits, only titles and abstracts were searched from the articles published till 31^st^ August 2020.[23, 24] A list of sample queries used for searching has been provided in **Table-1** with the number of hits observed.

**Table 1:** PubMed sample queries used in the present study and respective no. of hits obtained till August 2020.

| No. | Topic | PubMed Advanced queries | Total no. of records in PubMed |
| --- | --- | --- | --- |
| 1 | All abstracts | all[sb] AND hasabstract[text] | 20816630 |
| 2 | Stroke related abstracts | (stroke[TIAB] OR Cerebrovascular[TIAB]) AND (gene[TIAB] OR genes[TIAB]) AND hasabstract[text] | 9626 |
| 3 | Hemorrhagic Stroke related abstracts | ("Intracerebral hemorrhage"[TIAB] OR "Hemorrhagic Stroke"[TIAB] OR "Subarchanoid hemorrhage"[TIAB]) AND (gene[TIAB] OR genes[TIAB]) AND hasabstract[text] | 585 |
| 4 | Ischemic Stroke related abstracts | ("Ischemic Stroke"[TIAB] OR (TOAST[TIAB] AND (Classification[TIAB] OR Subtypes[TIAB]))) AND (gene[TIAB] OR genes[TIAB]) AND hasabstract[text] | 2738 |
| 5 | Large artery atherosclerosis | "Large artery atherosclerosis"[TIAB] AND (gene[TIAB] OR genes[TIAB]) AND hasabstract[text] | 103 |
| 6 | Small vessel disease | "Small vessel disease"[TIAB] AND (gene[TIAB] OR genes[TIAB]) AND hasabstract[text] | 288 |
| 7 | Cardioembolic disease | "Cardioembolic disease"[TIAB] AND (gene[TIAB] OR genes[TIAB]) AND hasabstract[text] | 117 |
| 8 | Other determined etiology | "Other determined etiology"[TIAB] AND (gene[TIAB] OR genes[TIAB]) AND hasabstract[text] | 3 |
| 9 | Undetermined etiology | "Undetermined etiology"[TIAB] AND (gene[TIAB] OR genes[TIAB]) AND hasabstract[text] | 45 |
| 10 | Intracerebral hemorrhage | "Intracerebral hemorrhage"[TIAB] AND (gene[TIAB] OR genes[TIAB]) AND hasabstract[text] | 424 |
| 11 | Subarchanoid hemorrhage | "Subarchanoid hemorrhage"[TIAB] AND (gene[TIAB] OR genes[TIAB]) AND hasabstract[text] | 434 |

### Model development

Document frequency (DF) related to a query is defined as the number of hits fetched by searching the database. DF can be easily normalized by the total no. of entries in the database. Similarly, pointwise mutual information (PMI) is another important metric often employed to find the association between two random variables (RV). A normalized form of PMI (nPMI) was derived by Bouma et al.[22] In several published works, nPMI were employed to estimate the association between entities like genomic repeats, stress, virulence, computational tools, drug-discovery related keywords, etc.[25, 26] Following a similar approach of association mining, individual and co-occurrences of each gene symbol (RV1) and stroke-related keywords (RV2) in PubMed titles and abstracts was calculated using normalized DF (nDF) which was further utilized to compute nPMI. This nPMI value represents the strength of association between the gene (genotype) and stroke (phenotype).

### Performance evaluation

Performance of the model was assessed using receiver operating characteristic (ROC) and precision-recall (PR) curve analysis on a cumulative dataset of human housekeeping (negative) [27] and already published stroke-related genes (positive). A positive gene set was constructed by compiling gene lists provided in the stroke-related research articles published during the last decade. [7, 18, 28] Different DF and nPMI value pairs were used to find the best model for stroke-associated gene identification.

### Pathway analysis

To achieve insight into the biological roles and pathological mechanisms of stroke and its etiologies, we examined the biological ways that significantly overlapped with the curated stroke and etiology gene sets. For this, we calculated common genes related to stroke etiology gene sets and the genes wiki-pathways [29] and executed statistical testing to measure the significance of the overlaps. To achieve insight into the biological functions and pathological mechanisms of stroke and its etiologies, we recognized the biological pathways that significantly enriched with the curated stroke and etiology gene sets. Using cluster Profiler R package,[30] enrichment analysis was performed on three major pathway databases namely KEGG (https://www.genome.jp/kegg/) (release 96.0),[31] WikiPathways (https://www.wikipathways.org/) (release September 2020)[29] and Reactome (https://reactome.org/) (version 75)[32] which are curated, comprehensive and rich data sources on human metabolic pathways. To initiate the analysis, probable stoke associated (PSA) gene symbols were first converted to ENTREZ ids and then pathway enrichment was done with a *p-value* cutoff of 0.05 and was adjusted by the Bonferroni method.[33]

## Results

### Genes associated with stroke and its types

PubMed advanced searched using stroke-related keywords as mentioned in the **Table-1** and associations were calculated using nPMI between each gene symbol and queries. To reduce the false hits, only titles and abstracts were searched from the articles published till 31^st^ August 2020.

A total of 2,785 (9.4%) genes were found to be linked to stroke risk. Based on stroke types, 1,287 (46.2%) and 376 (13.5%) genes were found to be associated with the risk of IS and HS respectively. Further stratification of IS based on TOAST classification, it was found that 86 (6.6%) genes were confined to Large artery atherosclerosis (LAA); 131 (10.1%) and 130 (10%) genes were related with the risk of small vessel disease (SVD) and Cardioembolism (CE) subtypes of IS. Circos diagram for the identified genes associated with stroke types and subtypes are represented in **Figure-1**.

**Figure 1:**
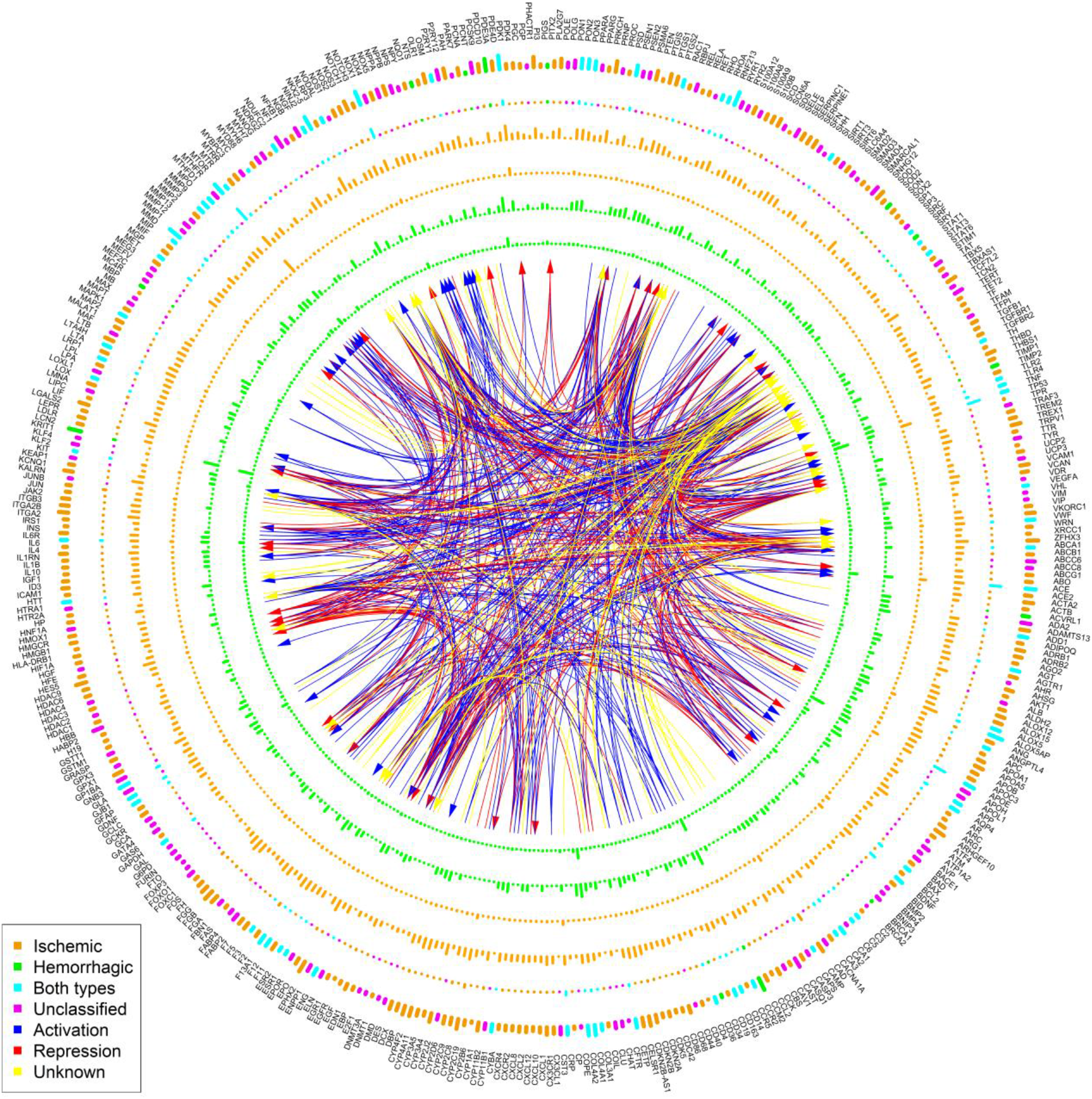
Prediction of the genes associated with stroke and its sub-types using normalized pointwise mutual information (nPMI) and document frequency (nDF) calculated from PubMed database. The outermost track of the circus plot represents the selected human gene symbols (#440) based on the nDF cut-off. The next two inner tracks with different colours show the nPMI and nDF values respectively; the colour codes are as follows: symbols associated with ischemic stroke: orange, hemorrhagic stroke: green, both stroke types: cyan and unclassified: magenta. nPMI and nDF values of the genes related with ischemic stroke have been presented in the next two orange inner tracks. Similarly, the two innermost green tracks displays nPMI and nDF values of the genes related with hemorrhagic stroke. The height of the bars indicates the nPMI and nDF values. The transcriptional regulatory links have been created between the transcription factors and pointing towards their target genes in different colours with codes: blue: activation, red: repression, yellow: unknown based on the manually curated database TRRUST (v2).

Total 28,281 human gene symbols (**Supplementary Table-T1**) were extracted from the NCBI gene database with the status tag “Active” and used for calculation of nPMI with query no. 2 from **Table-1**. 2,785 (9.8%) symbols were found to be associated (having positive or negative nPMI values) with stroke (set A). To determine the stroke subtypes, nPMI was computed with query no. 3 and query no. 4 (**Table-1**) for these 2,785 gene symbols (**Supplementary Figure-S1**) resulting 1,294 (46.5%) and 376 (13.5%) genes in association with IS and HS respectively. The rest of the symbols were marked as “Unclassified”. Further filtering using DF values (>5) and removing symbols (#11) like CAT, IMPACT, SET, etc. which are common English words, were listed 441 PSA genes along with their types (**Figure-1**). Gene symbols that were found to be in association with both types (HS and IS) of strokes were tagged as “Both Types” (**Figure-1**). Mining manually curated TRRUST database v2 (transcriptional regulatory relationships unraveled by sentence-based text-mining) (https://www.grnpedia.org/trrust/)[34] catalogued all the transcription factors and their target genes along with existing interaction types (activation, repression, or unknown) (**Figure-1**). A prognostic panel of 9 genes signature consisting of CYP4A11, ALOX5P, NOTCH, NINJ2, FGB, MTHFR, PDE4D, HDAC9, and ZHFX3 can be treated as a diagnostic marker to predict individuals who are at the risk of developing stroke with their subtypes

### Genes associated with stroke sub-types

Queries no. 5-9 from **Table-1** have been used in nPMI model for stratification of genes associated with IS subtypes as per TOAST classification. 131 (10.1%) genes were confined to Small Vessel Disease (SVD) followed by 130 (10%) Cardioembolism (CE), 86 (6.6%) Large artery atherosclerosis (LAA), 30 (2.3%) Undetermined etiology (UDE) and 7 (0.5%) Other determined etiology (ODE) (**Supplementary Figure-S2**). While classifying HS sub-types using queries no. 10-11, 292 (77.8%), and 132 (35.2%) were predicted as Intracerebral hemorrhage (ICH) and Subarachnoid hemorrhage (SAH) respectively (**Supplementary Figure-S3**). A subset of PSA (#140) genes associated with different stroke subtypes was represented in **Figure-2**. Complete lists along with DF and nPMI values can be found in **Supplementary Table T2-T10**.

**Figure 2:**
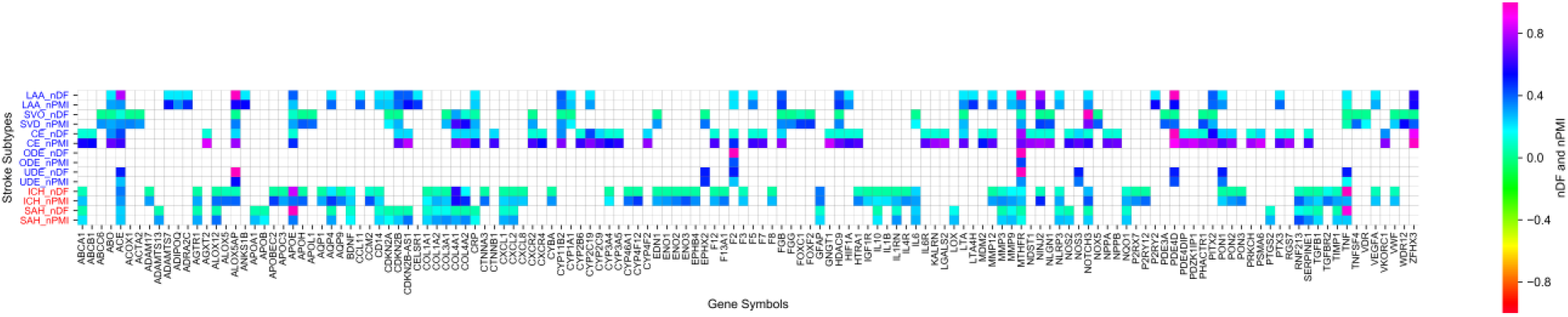
Prediction of genes associated with stroke types and subtypes. X-axis represents human gene symbols and Y-axis represents different stroke types and subtypes. Sub-types of Ischemic stroke (IS) are in blue and Hemorrhagic stroke (HS) are in red. Values of normalized Document frequency (nDF) (ranging 0 to 1) and normalized pointwise mutual information (nPMI) (ranging −1 to 1) are shown using color bar. Abbreviation of different stroke subtypes are as follows: SVD-Small Vessel Disease, CE-Cardioembolism, LAA-Large artery atherosclerosis, UDE-Undetermined etiology, ODE-Other determined etiology, ICH-Intracerebral hemorrhage SAH-Subarchanoid hemorrhage.

### Evaluation of nPMI model

For performance evaluation of the developed nPMI model, a list of already published 7,431 genes consisting of 2,168 (29%) stroke-related (positive set) and 5,263 (71%) human housekeeping genes (negative set) were constructed. An intersection of 1,144 genes was found between set A and published gene sets (positive: 581, negative: 563) which was used for evaluation. Multiple models were built using different DF (ranging from 1 to 5) and nPMI (ranging from 0.05 to 0.5) cut-offs and the best model having Accuracy: 0.64, Sensitivity: 0.63, Specificity: 0.65, and Precision: 0.66 was reported at DF cut-off 2 and nPMI cut-off 0.1. Performance evaluation by receiver operating characteristic (ROC) and Precision-Recall (PR) curves analysis resulted Area under curve (AUC) values of 0.64 and 0.63 respectively (**Figure-3**).

**Figure 3:**
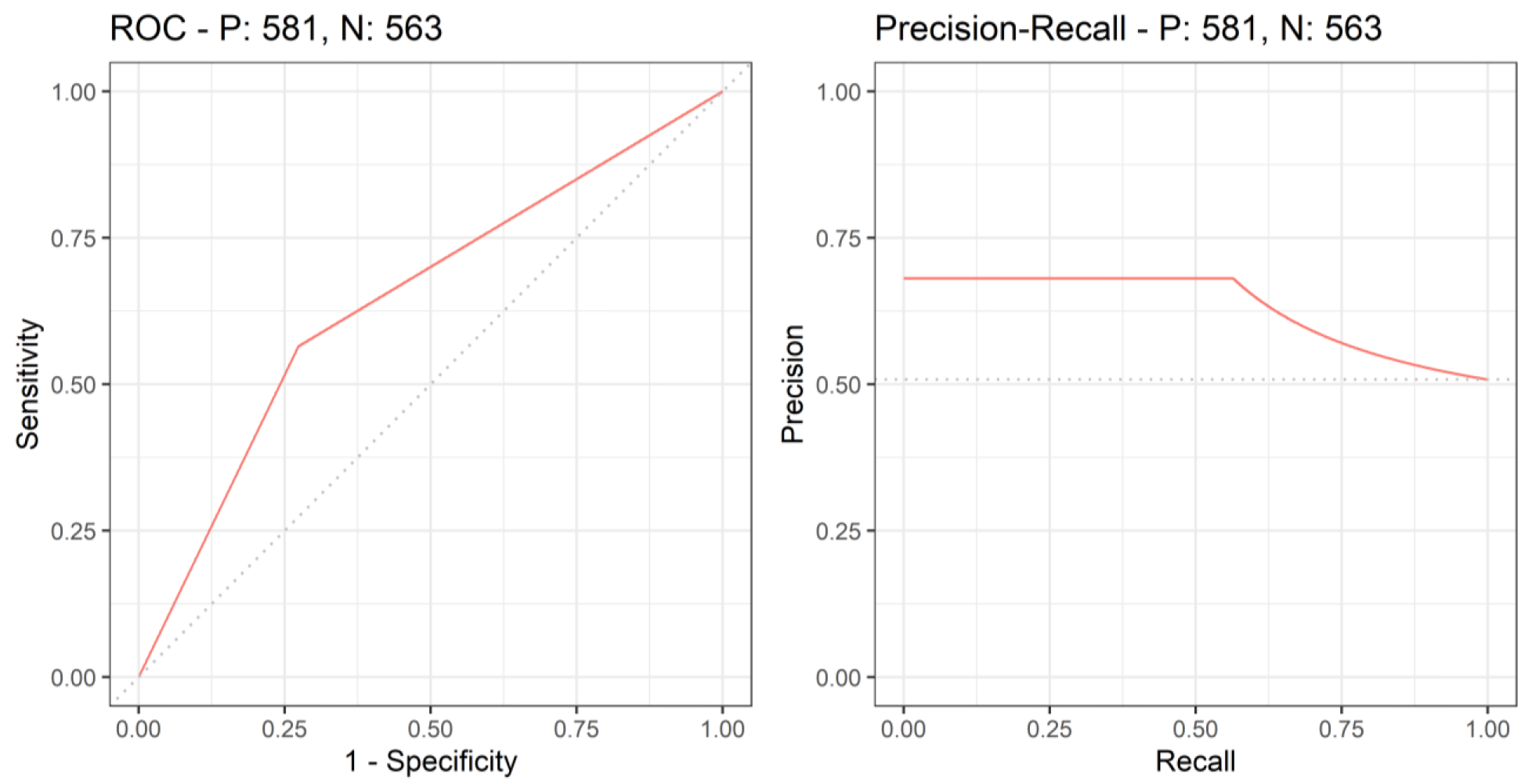
Receiver operating characteristic (ROC) and Precision-Recall (PR) curve analysis for the evaluation of developed normalized pointwise mutual information (nPMI) based model for classification of stroke related genes. Using different normalized document frequency (nDF) and nPMI cut-offs ROC and PR curves have been plotted. Area under curve (AUC) values for ROC and PR curves are 0.64 and 0.63 respectively.

### Stroke can influence many pathways

To reduce false-positive hits, pathway enrichment analysis was done using 190 genes (**Supplementary Table-T11**), a subset of PSA gene symbols that were manually curated and already known to be associated with stroke. Analysis with Reactome, WikiPathways, and KEGG resulting 53, 32, and 35 unique pathways enriched with aforementioned stroke-associated genes (**Supplementary Table-T12, Table-T13, and Table-T14**). However manual curation of these lists of pathways showed promising results with WikiPathways and Reactome which were presented in **Figure-4** and **Supplementary Figure-S4**. Findings for the genes associated with biological Network pathways including Kegg and Reactome are reported in **Figure-4**.

**Figure 4:**
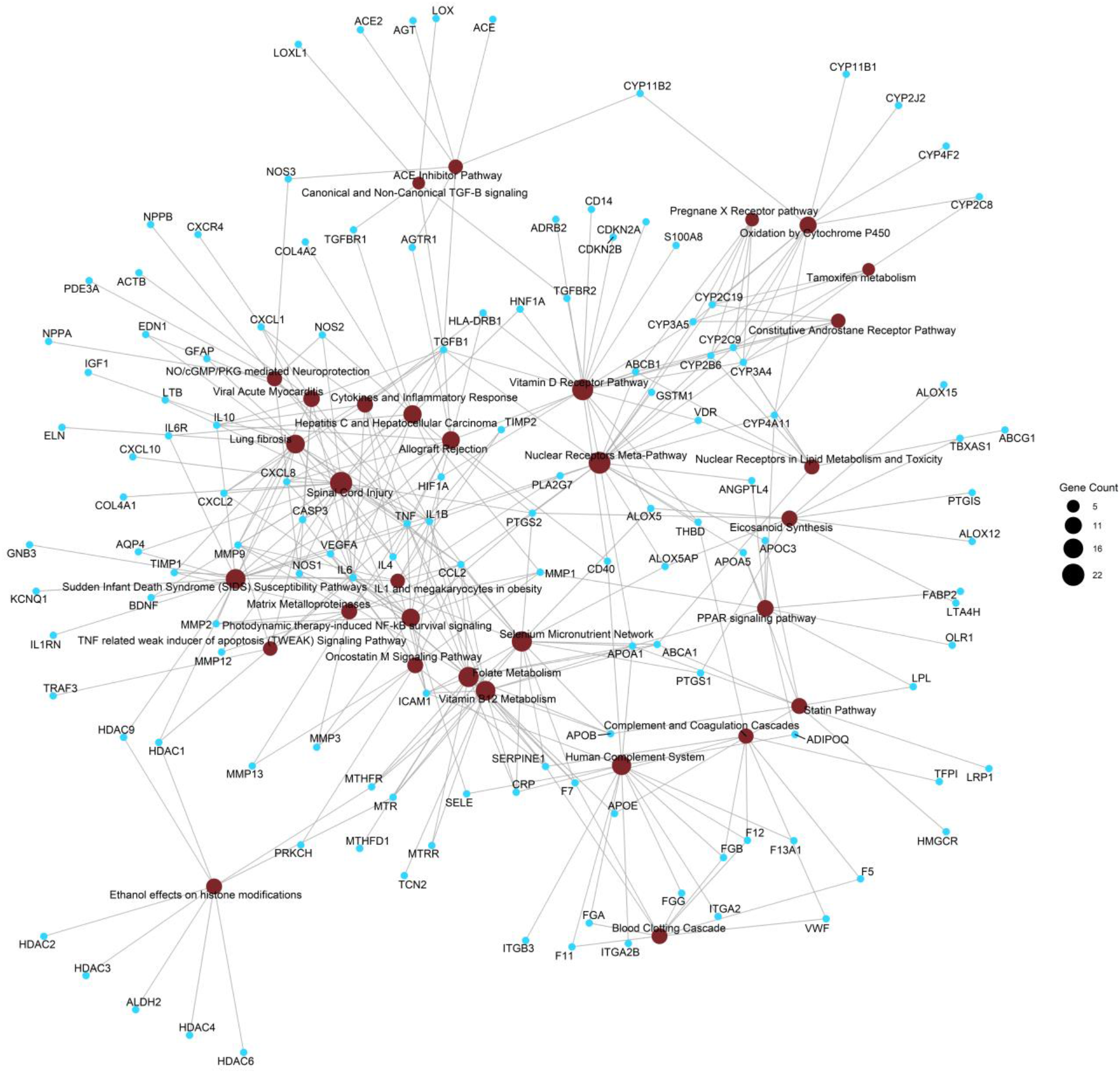
Enrichment analysis for finding the important pathways associated with the predicted stroke genes. Analysis has been performed using WikiPathways database. Brown and cyan nodes are representing pathways and genes respectively. No. of genes associated with a pathway is represented by size of the brown node. Connecting links are in grey color.

## Discussion

Emerging evidences from published meta-analysis and GWAS studies suggests that several genetic variants (MTHFR, MMP9, PDE4D, CYP4A11, ALOX5P, NOTCH, NINJ2, FGB, eNOS, PITX2, ZFHX3, HDAC9, ABO, etc) have been identified, even though the extent of the effect of each variant is regarded as inter-varying within different populations including Asian, Caucasian, African.[1, 1, 7, 14, 35–39, 39–44] The findings, however, are often unclear and hard to interpret. Genetic association studies in diverse stroke populations negate matters of the restricted patient population by the a priori choice of a functionally relevant gene and its relation with a specific phenotype. Improving patient outcomes in stroke requires a rapid and accurate prediction of stroke and its subtypes. The genetic signature could help it to distinguish or calculate the incidence of hemorrhagic and ischemic stroke along with its subtypes and may also help in the prognosis of further risk of stroke recurrence.

Our findings are confined specifically to predict the associated genes for the predisposition of ischemic or hemorrhagic stroke. A prognostic panel of 9 genes signature consisting of CYP4A11, ALOX5P, NOTCH, NINJ2, FGB, MTHFR, PDE4D, HDAC9, and ZHFX3 can be designed and treated as a diagnostic marker to predict individuals who are at the risk of developing stroke with their subtypes. Developing this genetic markers panel seems to offer hope of significantly better sensitivity and specificity may provide a quick and reliable assessment with revolutionize stroke management. It will also reduce cost and timing for preventing the stroke incidence in susceptible individuals having a chance of developing stroke with LVD, SVD and CE subtypes and HS. These genetic markers could also have the potential to enter into a routine clinical use despite their obvious promise by only single validation of our findings.

## Limitations

The current study has certain limitations in spite of these interesting results. Meanwhile the datasets fused were from the published studies in patients with stroke, misdiagnosis or misclassification of stroke subtypes could have potentially influenced our findings. Also, we could not access the original SNP genotype data, we had to use summary data from published stroke GWAS and candidate gene studies, which prohibited us to address the common genetics of complex traits and could have affected our results. Moreover, as the various testing corrections we used in our statistical analyses may be inadequate to clarify all biases, permutation testing should be used to adjust the results at the single SNP level. Furthermore, we needed transcriptomic and epigenetic data, which may contribute to the identification of additional potential causal mechanisms and links. False positives are highly expected since the method was predicting the genes using PubMed title and abstracts. However, false positives in our findings were rectified using manual curation.

## Conclusion

A novel text data-driven method was developed to identify genes associated with stroke types and their subtypes. Our findings might offer certain directive implications for further exploring the diagnostic and prognostic genetic markers to empower the molecular targeting treatment for stroke prevention.

## Supporting information

GD_stroke_supplementary_13092021

GD_stroke_supplementary_13092021

## Notes

**Conflict of Interest** No potential conflict of interest

**Funding Source** None

### Competing Interest Statement

The authors have declared no competing interest.

