## Supplementary material for "Potential Key Genes Associated with Stroke types and its subtypes: A Computational Approach": GD_stroke_supplementary_13092021

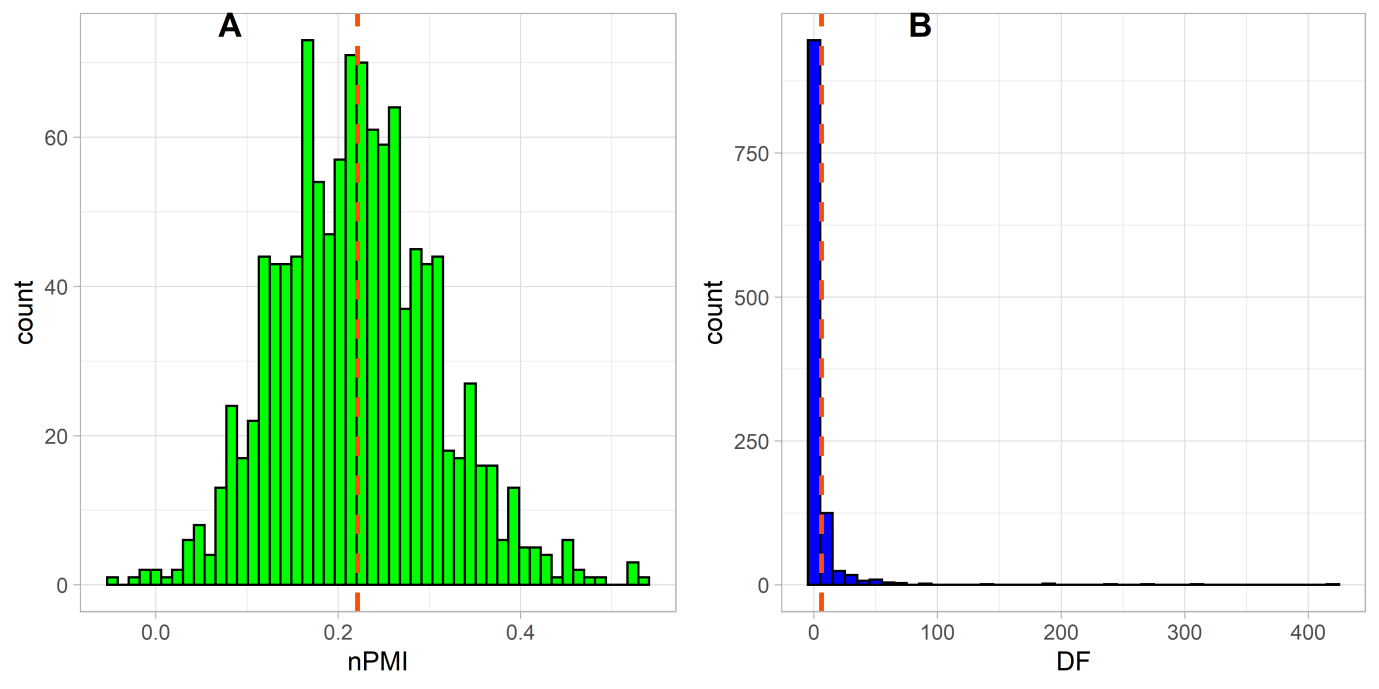

Figure S1: Distributions normalized pointwise mutual information (nPMI) (A) and document frequency (DF) (B) for the 2,785 genes predicted to be associated with stroke. Y-axis represents the counts of gene symbols. The position of mean is shown in the red dotted line.

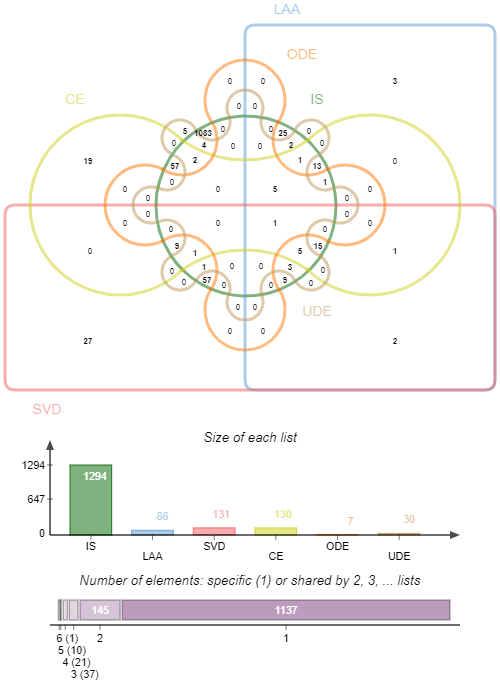

Figure S2: Venn diagram showing common genes among Ischemic stroke (IS) type (green) and its subtypes i.e. Large artery atherosclerosis (LAA) (blue), Small vessel disease (SVD) (red), Cardioembolic disease (CE) (yellow), Other determined etiology (ODE) (orange) and Undetermined etiology (UDE) (brown) as predicted using nPMI model and different queries (no. 3, 10 and 11) from Table 1.

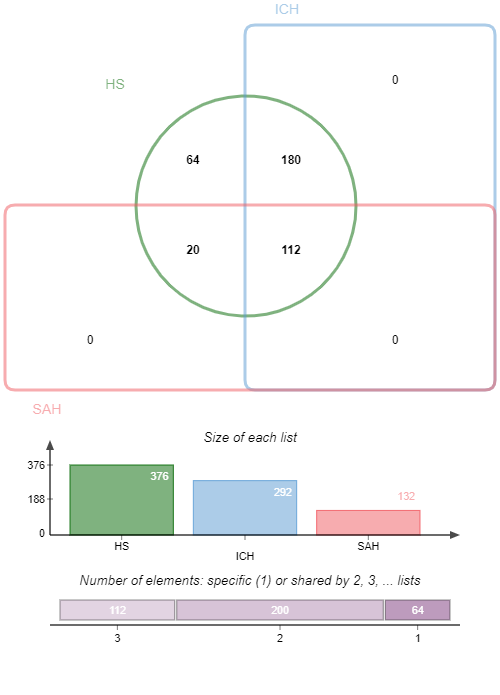

Figure S3: Venn diagram showing common genes among hemorrhagic stroke (HS) type (green) and its subtypes i.e. Subarachnoid hemorrhage (SAH) (red), Intracerebral hemorrhage (ICH) (blue) as predicted using nPMI model and different queries (no. 3, 10 and 11) from Table-1.

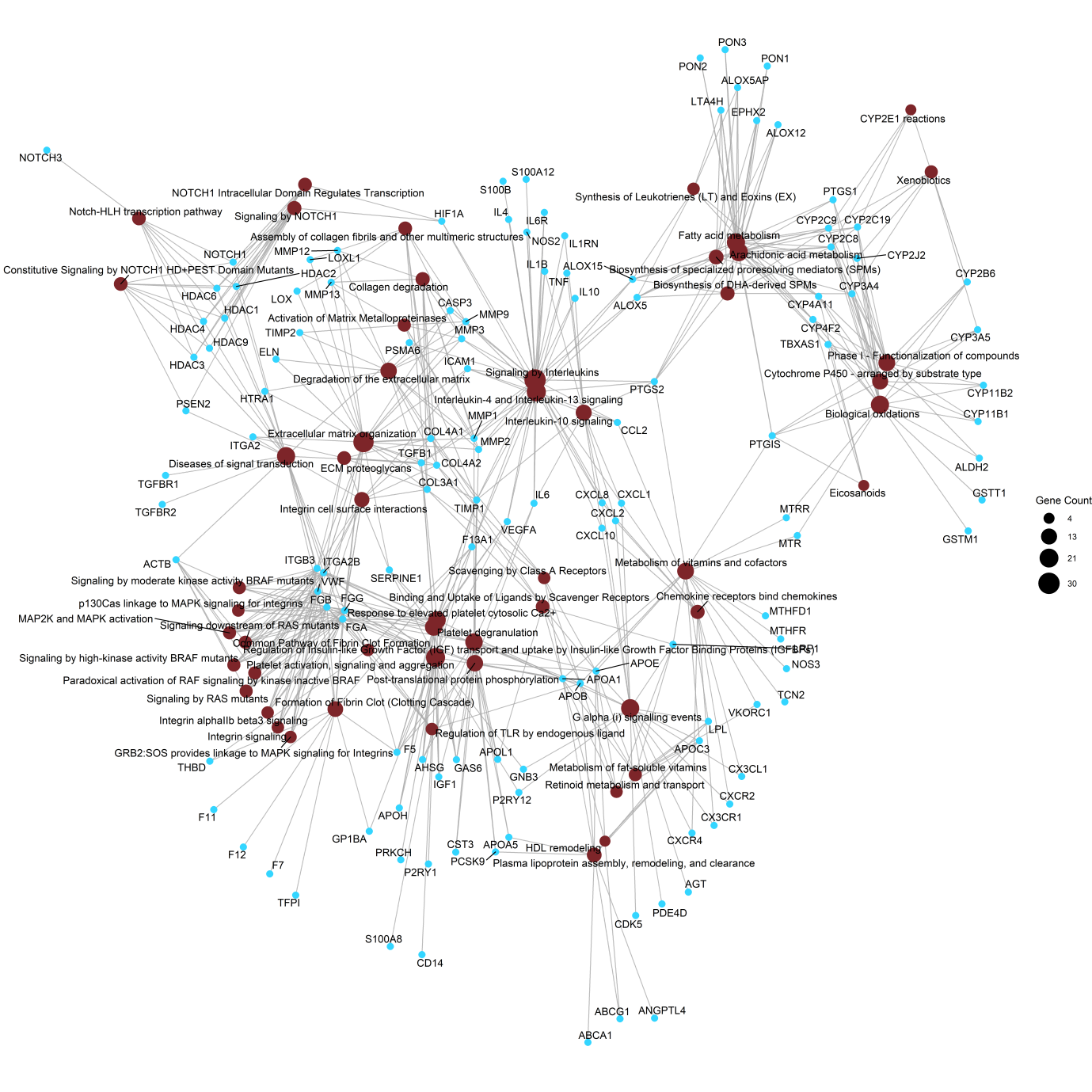

Figure S4: Enrichment analysis for finding the important pathways associated with the predicted stroke genes. Analysis has been performed using the Reactome database. Brown and cyan nodes are representing pathways and genes respectively. No. of genes associated with a pathway is represented by the size of the brown node. Connecting links are in grey.
